# Macrolactin A Is an Inhibitor of Protein Biosynthesis in Bacteria

**DOI:** 10.1101/2024.02.10.575802

**Authors:** Alexey S. Vasilchenko, Dmitry A. Lukyanov, Diana S. Dilbaryan, Konstantin S. Usachev, Darya V. Poshvina, Arina A. Nikandrova, Arina N. Imamutdinova, Natalia S. Garaeva, Aydar G. Bikmullin, Evelina A. Klochkova, Amir Taldaev, Alexander L. Rusanov, Daniil D. Romashin, Natalia G. Luzgina, Ilya A. Osterman, Petr V. Sergiev, Anastasia V. Teslya

## Abstract

The macrolide antibiotic, macrolactin A (McA), has been known for its antimicrobial properties since the late 1980s, but the mechanism of its antibacterial activity is still unknown. In this study, we investigated the microbiological and molecular characteristics of McA antimicrobial activity. McA effect on bacteria was found to be both bacteriostatic and bactericidal, depending on species and strains. Regarding the mechanism of action of McA, the following important results were obtained: 1) using *in vivo* and *in vitro* systems, we showed that McA is an inhibitor of protein synthesis in bacteria; 2) the concentration of McA required to inhibit protein synthesis in the *E. coli* cell-free model was found to be 50 times lower than the concentration required in the *S. aureus* cell-free model; 3) the toe-printing assay revealed that McA inhibits the first step of elongation stage of protein synthesis; 4) we identified single and multiple nucleotide polymorphisms in the gene encoding the translation elongation factor Tu (EF-Tu) by annotating the genomes of McA-resistant *Bacillus pumilus* McA^R^ and its parental strain. Molecular modeling showed that the McA molecule can form non-covalent bonds with amino acids at the interface of domains 1 and 2 of EF-Tu, characterized by a relatively high docking score. Overall, our study demonstrated that McA acts as an elfamycin-like antibiotic (targeting EF-Tu), addressing a substantial gap in our understanding of the mechanism of action of macrolactin A, a representative member of macrolides.

## 1. INTRODUCTION

Analyzing the chronology of the discovery of new antibiotics, it becomes clear that by the middle of the 20th century, the range of antibiotics had acquired its modern form. Most antibiotics discovered in the last 30 years demonstrate activity mainly against Gram-positive bacteria (Lewis, 2020), while the World Health Organization (WHO) has declared Gram-negative bacteria of the *Enterobacteriaceae* family as a priority in terms of antibiotic resistance (Tacconelli et al., 2018).

Certain macrolide antibiotics have been used safely and effectively to treat bacterial pneumonia, gonorrhea, and other diseases over many years of use (Patel, Hashmi, 2023). Erythromycin, first extracted from the soil bacterium *Saccharopolyspora erythraea*, is the prototypical macrolide antibiotic (McGuire et al. 1952). This class of antibiotics is characterized by a macrolactone ring with one or more attached sugars (Myers et al., 2021). The macrolide class includes macrolactins, which have a predominantly 24-membered lactone ring in their structure. The first members of macrolactins, including macrolactin A (McA), were described in 1989 as secondary metabolites of an unidentified microorganism isolated from deep sea sediments of the North Pacific Ocean (Gustafson et al. 1989). Today, 33 different macrolactins have been isolated and structurally described (Wu et al. 2021). Macrolactins are synthesized non-ribosomally using polyketide synthetases, which are widely distributed in microorganisms inhabiting both marine and terrestrial environments (Zhang et al. 2022).

Since their discovery, antibacterial properties of macrolactins have not been extensively studied. The spectrum of targeted bacterial species is limited to known human pathogens such as *S. aureus* and *E. coli* and several phytopathogens (Gustafson et al. 1989; Han et al. 2005; Kim et al. 2011; Yuan et al. 2012). The mechanism of action of macrolactins is still poorly understood.

Existing studies describe the features of the mechanism of action of certain macrolactins. For instance, Zotchev et al. (2006) utilized the PASS/PharmaExpert software algorithm to predict that the antibacterial activity of macrolactins could be associated with the inhibition of the bacterial H^+^-transporting two-sector ATPase. The study by Yoo et al. (2006), showed the inhibitory effect of macrolactin N on bacterial peptide deformylase. Sohn et al. (2008) demonstrated that macrolactin S and macrolactin B have selective inhibitory activity against NADPH-dependent β-ketoacyl-ACP reductase (FabG), an enzyme involved in fatty acid synthesis in staphylococci. In the study by Chen et al. (2020), *Agrobacterium tumefaciens* cells treated with macrolactin A were imaged using transmission electron microscopy, which revealed undivided cells with evidence of septal inhibition.

However, none of these studies aimed to test the hypothesis that the target of macrolactins, like other macrolide antibiotics, is protein synthesis. In our study, we unraveled the mechanism of action of McA on bacterial cells. We characterized the qualitative and quantitative features of the antibacterial action of McA on bacteria with different structural organization of the cell wall. The effect of McA on protein biosynthesis was tested and described in detail using various bioreporter systems, as well as the analysis of genomic mutations of the resulting McA- resistant bacterial strain. We discovered a probable target for McA in the protein biosynthesis machinery, which is the elongation factor EF-Tu. Molecular modeling revealed that the potential binding site of McA distinguishes it from known EF-Tu inhibitors such as kirromycin and pulvomycin.

## 2. RESULTS AND DiSCUSSION

### 2.1 Identification of the chemical structure of macrolactin A

The chemical structure of the purified antibacterial metabolite was established using LC/MS and NMR. Analysis of the mass spectrum (Figure S1) showed that the antimicrobial fraction corresponds to the molecular ion [M+Na]^+^ 425.06794, for which the brutto formula C_24_H_34_O_5_ was calculated. The MS result was supported by the the following NMR data:^1^H NMR (700 MHz, methanol-d4): (ppm) 5.56 (2-H), 6.63 (3-H), 7.24 (4-H), 6.19 (5-H), 2.44 (6-H2), 4.28 (7-H), 5.77 (8-H), 6.59 (9-H), 6.14 (10-H), 5.56 (11-H), 2.54 (12-Ha), 2.34 (12-Hb), 3.89 (13-H), 1.62 (14-H2), 4.34 (15-H), 5.58 (16-H), 6.20 (17-H), 6.09 (18-H), 5.68 (19-H), 2.22 (20-Ha), 2.14 (20-Hb), 1.52 (21-H2), 1.72 (22-Ha), 1.67 (22-Hb), 5.03 (23-H), 1.28 (24-H3).

13C NMR (176 MHz, methanol-d4): (ppm) 169.25 (C-1), 119.22 (C-2), 146.69 (C-3), 132.23 (C-4), 132.50 (C-5), 44.10 (C-6), 73.61 (C-7), 138.81 (C-8), 127.24 (C-9), 132.73 (C-10), 129.65 (C-11), 37.81 (C-12), 70.47 (C-13), 45.16 (C-14), 71.07 (C-15), 136.48 (C-16), 132.56 (C-17), 132.97 (C-18), 136.43 (C-19), 34.31 (C- 20), 27.08 (C-21), 37.16 (C-22), 73.55 (C-23), 21.41 (C-24).

A direct comparison of the 1H and 13C NMR data of the antimicrobial sample in MeOH-d6 with the previous reports shown almost identical values of chemical shifts with the McA produced by the *Bacillus subtilis* strain DSM 16696 (Romero-Tabarez et al. 2006).

### 2.2 Antimicrobial spectra of macrolactin A

The first study on the antibacterial activity of McA was conducted in 1989 and showed an antibiotic effect on Gram-positive *Staphylococcus aureus* and *Bacillus subtilis* at concentrations of 5 and 20 μg/disk, respectively (Gustafson et al., 1989). Subsequent studies have shown that McA is capable of inhibiting the growth of *S. aureus* IFO 12732 and *B. subtilis* IFO 3134 at concentrations of 10 and 60 μg/mL, respectively, while not inhibiting the growth of *E. coli* IFO 3301 and *Salinivibrio costicola* ATCC 22508 (Nagao et al., 2001). In the study by Romero-Tabarez et al (2006), McA used at a concentration of 50 μg/disk, inhibited the growth of various strains of *S. aureus* with inhibition zones ranging from 18 to 35 mm. However, it showed no activity against different strains of *Enterococcus faecalis*.

In the present study, the qualitative parameters of McA activity were evaluated using a panel of test strains comprising a wide range of both Gram-positive and Gram-negative bacteria (Table 1). It was found that there was no clear correlation between MBC levels and the type of bacterial cell wall. Among the test strains, the highest sensitivity to the activity of McA was observed in strains belonging to *Pectobacterium* genera. Interestingly, in the study of Maung et al. (2022), McA did not inhibit the growth of *P. carotovorum* even at a concentration of 200 µg/ml.

**Table 1.** Spectrum of antimicrobial activity of macrolactin A determined as a minimum bactericidal concentration (MBC)

| Cell wall type | Microorganisms | MBC (McA), µg/mL |
| --- | --- | --- |
| Gram-positive | *Staphylococcus aureus* FDA209P | 150 |
|  | *Staphylococcus aureus* ATCC 29923 | >150 |
|  | *Staphylococcus aureus* ATCC 43200 | 150 |
|  | *Staphylococcus epidermidis* UTMN | 5 |
|  | *Listeria monocytogenes* EGDe | 5 |
|  | *Lactobacillus fermentum* | >150 |
|  | *Enterococcus faecium* 79OSAU | 75 |
|  | *Enterococcus faecium* UTMN | 75 |
|  | *Bacillus cereus* IP 5832 | 50 |
|  | *Bacillus pumilus* UTMN | 5 |
| Gram-negative | *Escherichia coli* ATCC 25922 | 37 |
|  | *Escherichia coli* K12 | >150 |
|  | *Pseudomonas aeruginosa* ATCC 287531 | 60 |
|  | *Pseudomonas brassicacearum* RCAM05391 | >150 |
|  | *Chromobacterium substugae* ATCC 31532 | 75 |
|  | *Xanthomonas campestris* | >150 |
|  | *Pectobacterium carotovorum* VKM B-1247 | 20 |
|  | *Pectobacterium versatile* VKM B-3416 | 10 |
|  | *Pectobacterium aquaticum* VKM B-3417 | 50 |

Among the Gram-positive bacterial strains, *B. pumilus, L.monocytogenes* and *S.epidermidis* exhibited the highest sensitivity to the McA (Table 1). McA did not exhibit bactericidal activity against any of the *S. aureus* strains tested in the experiment. However, the analysis of bacterial growth curves showed that McA had a bacteriostatic effect on this species.

### 2.3 Macrolactin A kills bacteria within two hours

The dynamics of bactericidal effect development (time-kill curves) upon exposure to McA was evaluated on the sensitive strain of *P. carotovorum* VKM B-1247. Time-kill curves are commonly used to assess the biocidal potential of various antimicrobial agents. The action of McA, taken at a concentration of 1 MIC, was bacteriostatic for the first four hours and became bactericidal after 8 hours of co-incubation (Figure 1a). A rapid bactericidal effect within 2 hours was achieved at a concentration of McA corresponding to 2 MIC (Figure 1a). Azithromycin demonstrated a similar dynamic of bactericidal and bacteriostatic effects (Figure 2 b).

**Figure 1.**
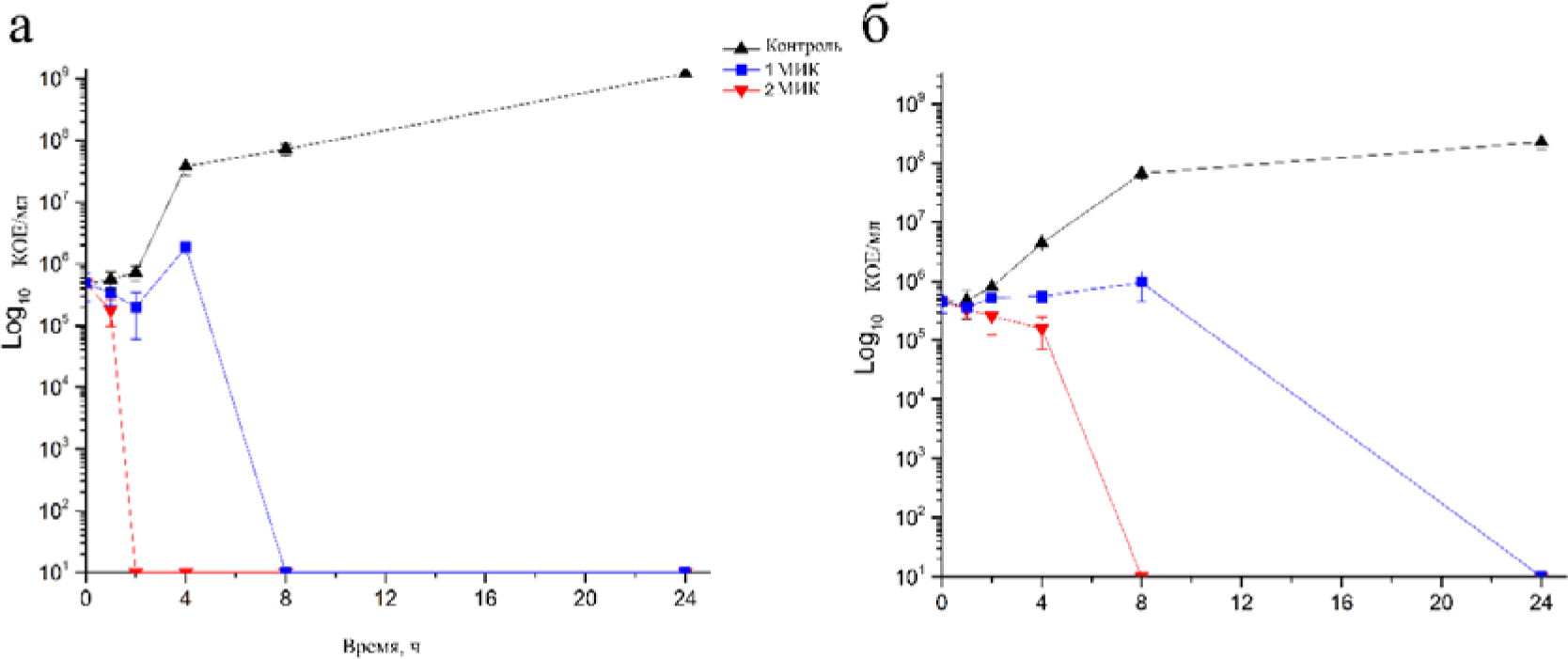
Dynamics of bactericidal effectiveness of macrolactin A (a) and azithromycin (b) against *P. carotovorum* VKM B-1247.The bacteria were co-incubated for 24 hours with antibiotics taken at concentrations corresponding to 1 MIC and 2 MIC.

**Figure 2.**
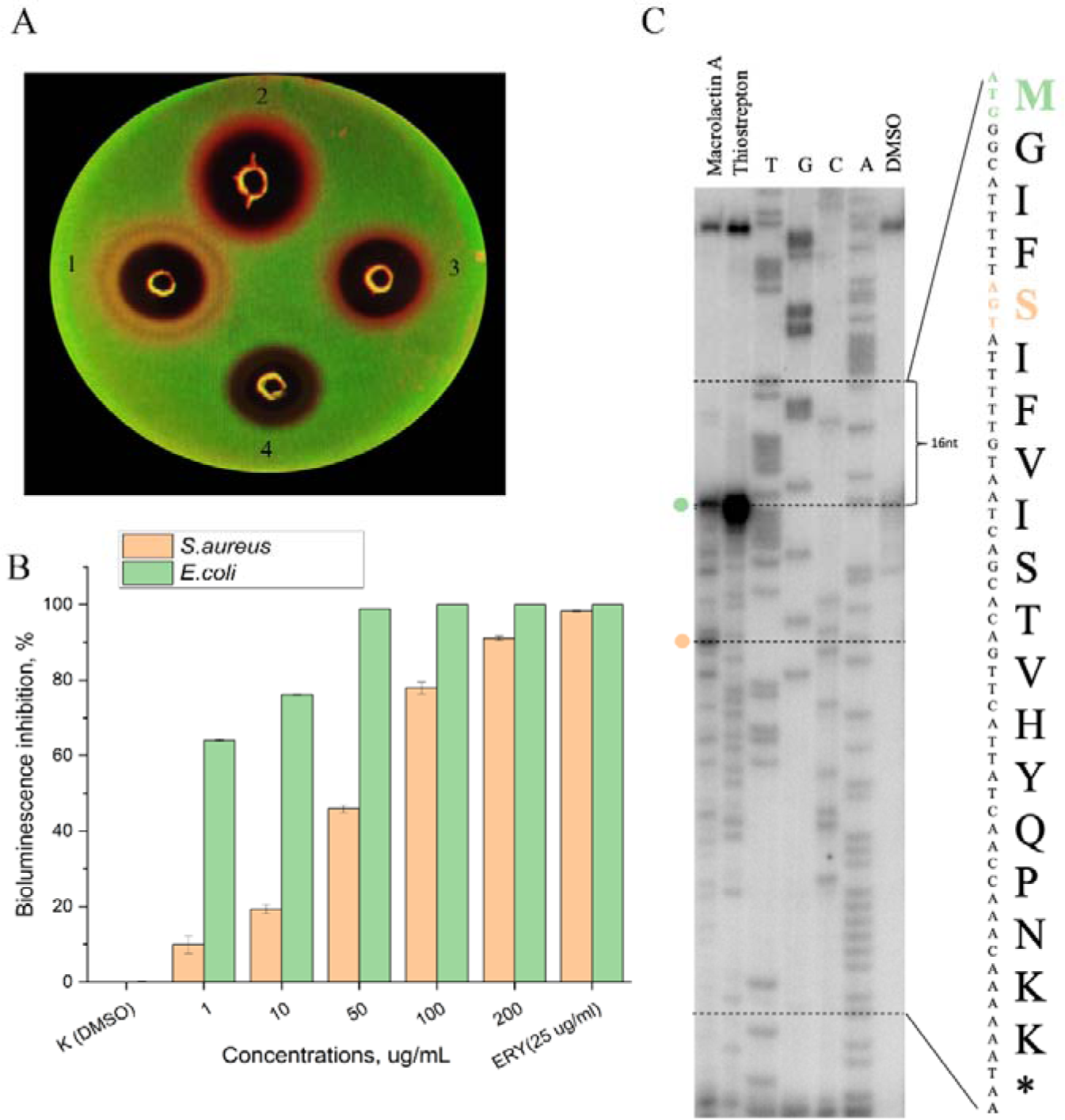
Macrolactin A is an inhibitor of ribosome translocation. A) Fluorescence detection of the reporter strain *E. coli* JW5503 ΔtolC transformed with the pDualrep2 plasmid when exposed to: 1) azithromycin, 2) tetracycline, 3) macrolactin A, 4) gentamicin. Images are constructed with an overlay of two detection channels where Katushka2S is displayed in red. B) Inhibition of protein synthesis by increasing concentrations of McA in *in vitro* cell- free translation in *E. coli* S30 and *S.aureus* RN4220 extracts. C) The scheme of toe-printing assay on the ermCL template: lane 1 corresponds to the *in vitro* cell-free translation system supplemented with 100 μg/mL of McA; lane 2 is thiostrepton taken for 50 μM (inhibits translation at the start codon (Orelle et al. 2013)); T, G, A, C are the lanes corresponding to sequencing reactions with serial stops at the corresponding nucleotides; lane 3 corresponds to the negative control (1% DMSO).

It is possible that some time is required for a sufficient amount of antibiotic to penetrate into the cell to develop an irreversible bactericidal effect. A static effect was observed during the first 8 hours, while the bactericidal effect was established after 24 hours of co-incubation (Figure 2a). According to existing studies, azithromycin concentration corresponding to MIC exert a bacteriostatic activity (Belanger et al. (2020)), while a bactericidal effect against Gram-negative bacteria (Elkashif et al., 2020; Singkham-In et al., 2021) and Gram-positive bacteria (Lemaire et al., 2009) was observed at concentrations exceeding the MIC value.

### 2.4 Macrolactin A inhibits bacterial protein synthesis *in vivo* and *in vitro*

A preliminary investigation of the mechanism of action of McA was conducted using the biosensor strain *E. coli* JW5503 ΔtolC pDualrep2. Using this reporter strain, an induction signal of Katushka2S was obtained from positive controls, namely azithromycin, tetracycline and gentamicin. In the case of McA, a fluorescent signal was detected and showed intensity comparable to that of tetracycline (Figure 2 A). In the study by Osterman et al. (2016), it was explained that aminoglycosides, including gentamicin, showed very weak or no induction of the Katushka2S reporter. One possible reason for such a response could be that the main mechanism of action of aminoglycosides is the induction of reading errors rather than ribosome stalling.

We confirmed the protein synthesis inhibitory activity of McA using *in vitro* assays recording luciferase mRNA translation (*E. coli* S30 extract) and sfGFP fluorescence (*S.aureus* S30 extract). It was found that the reduction of luciferase mRNA translation *in vitro* was observed in a concentration-dependent manner (Figure 2 B). McA significantly inhibited (IC50) the luciferase signal starting from the concentration of 1 μg/mL in the reaction mix reproducing *E.coli* model. Concentrations ≥ 50 μg/mL are required for a complete (IC 100) blocking of protein synthesis. At the same time, the biosynthetic machinery of cell-free model representing *S. aureus* exhibited a higher stability towards McA. The concentration of 1 μg/mL inhibited bioluminescence by only 10%, while the concentration of 50 μg/mL was found to be the IC50 (Figure 2 B).

### 2.5. Macrolactin A affects the elongation stage of protein synthesis

In a typical experiment, we compared the sites of ribosome stalling caused by the novel and the known translation inhibitors. The first nucleotide of the blocked ribosome P site on the mRNA and the last synthesized cDNA nucleotide are 16 nucleotides apart. It is convenient to use the thiostrepton antibiotic for comparison, as it is known to induce ribosome stalling at the first translation step, just when the start codon AUG is located at the ribosomal P site. Based on these data, we performed calculations for the codons located at the P site at the time of ribosome stalling (Figure 2 C). These codons were 1-AUG (M) and 5-AGT (S). We can see that the ribosome stalls at the 5th codon. Therefore, it can be hypothesized that this translation inhibitor McA can affect the elongation stage of protein synthesis.

We compared the sites of ribosome stalling caused by the novel and already known translation inhibitors. The first nucleotide of the P site of the ribosome blocked on mRNA and the last synthesized cDNA nucleotide are 16 nucleotides apart. It is convenient to use the thiostrepton antibiotic for comparison, as it is known to induce ribosome stalling at the first translation step, right when the start codon AUG resides at the ribosomal P site. Based on these data, we calculated the codons residing at the P site at the instant of ribosomal stalling (Figure 2 C). These codons were 1-AUG (M) and 5-AGT (S). We can see that ribosome stalls at the 5^th^ codon. Therefore, a hypothesis can be put forward that this translation inhibitor McA can affect the elongation stage of protein synthesis.

To confirm this hypothesis, we carried out selection of the macrolactin A-resistant clone of *B. pumilus.* We then compared the genomes of the resistant clone *B. pumilus* McA^R^ with those of the parental strain and found single and multi-nucleotide polymorphisms in the *tuf* gene, which encodes the translation elongation factor Tu (EF-Tu). The mutation in the *tuf* gene causes the substitution of Arg125 by His125 and Val197 by Ala197 (Figure 3, Table 2).

**Figure 3.**
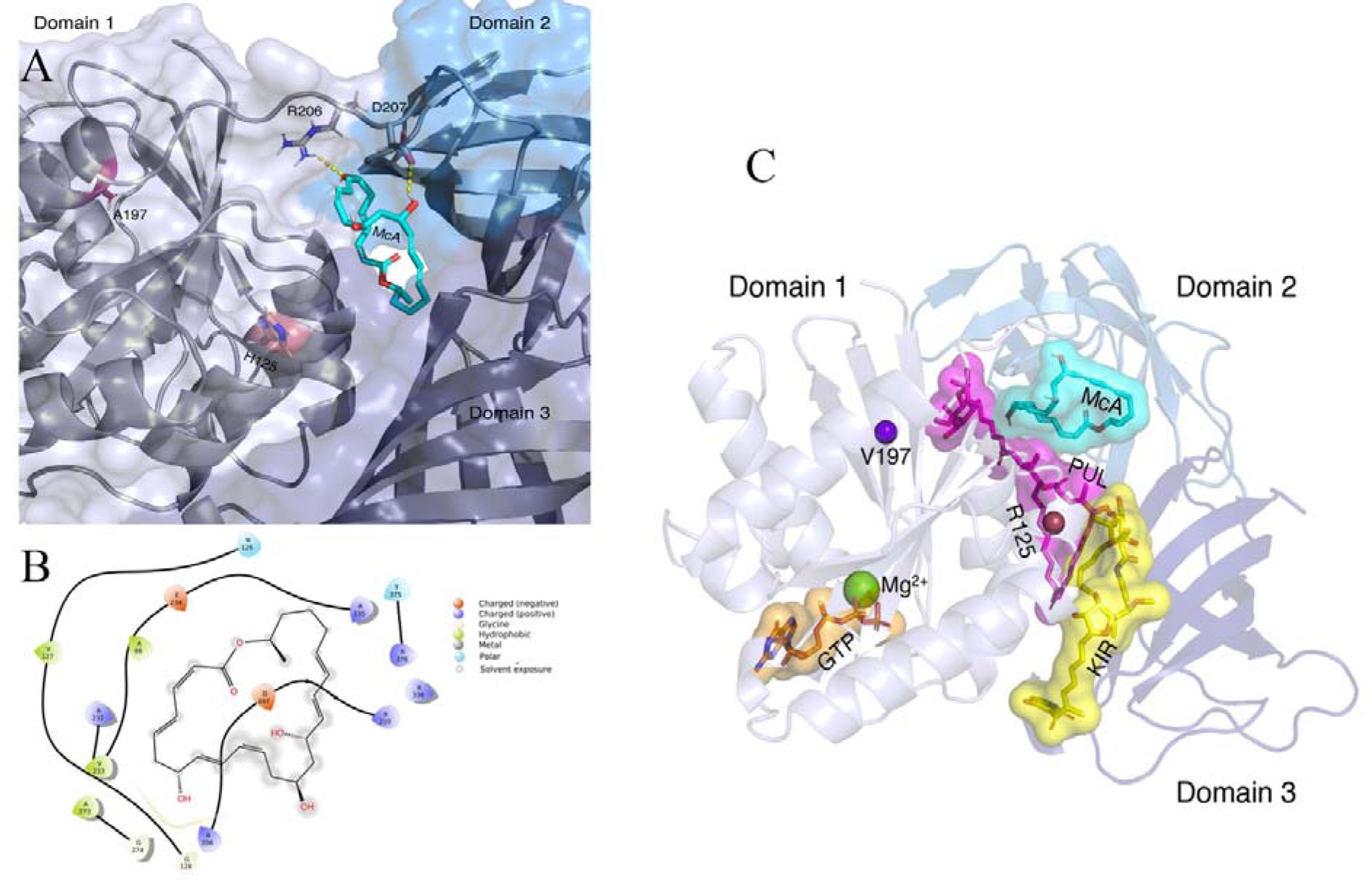
Elongation factor Tu is a central part of the bacterial translation machinery. (А) The structure of *B. pumilus* in an active (GTP-binding) conformation that is obtained via homology modeling. Enlarged fragment of the D1-D2 interface, where, according to molecular docking, there is a probable McA binding site (the best pose shown). Mutated and important interacted amino acids residues shown as sticks; important electrostatics bonds marked as yellow dashed lines. B) Predicted binding mode 2D diagram of McA and protein interactions. C) The general view on elfamycins interaction sites in EF-Tu. Kirromycin (KIR) binds to the D1-D3 interface EF-Tu (Warias et al. 2021), pulvomycin (PUL) has binding site extending from the domain 1-3 interface to domain 2, overlapping domain 1-2-3 junction (Parmeggiani et al. 2006).

**Table 2.** Mutations in the genome of *B. pumilus* McA-resistant strain.

| Nucleotide position in *B. pumilus* McA<sup>S</sup> | Type of mutation | *B. pumilus* McA<sup>S</sup> | *B. pumilus* McA<sup>R</sup> | Nucleotide position in the gene | Amino acid position | Effect |
| --- | --- | --- | --- | --- | --- | --- |
| 3,618,113 | SNP | A | G | 374/1191 | 125/397 | missense variant c.374 A>G p. Arg125His |
| 3,618,329 | SNP | CC | TT | 590-591/1191 | 197/397 | missense variant c.590-591 CC>TT p. Val197Ala |

Today, more than 30 antibiotics are known (kirromycin, enacylocin IIa, pulvomycin, GE2270 A, KKL-55) that target the prokaryotic elongation factor Tu and are therefore called elfamycins (Prezioso et al. 2017). Thus, the data obtained make it possible to replenish the list of known elfamycins with McA.

We used molecular docking to calculate the likely binding site of McA to EF-Tu. The calculations showed that McA probably binds to the D1-D2 interface EF-Tu (Figure 4 A); that is similar to the pulvomycin binding mode, where the binding interface is also at D1-D2 of EF-Tu. According to these data, we suppose the McA mechanism of activity is to prevent the formation of the ternary complex EF-Tu –GTP – aa-tRNA. The result of molecular docking showed that the McA molecule can form two significant non-covalent bonds with amino acid residue side chains’ Arg206 and Asp207 in the D1-D2 interface (Figure 4 B and 4C).

**Figure 4.**
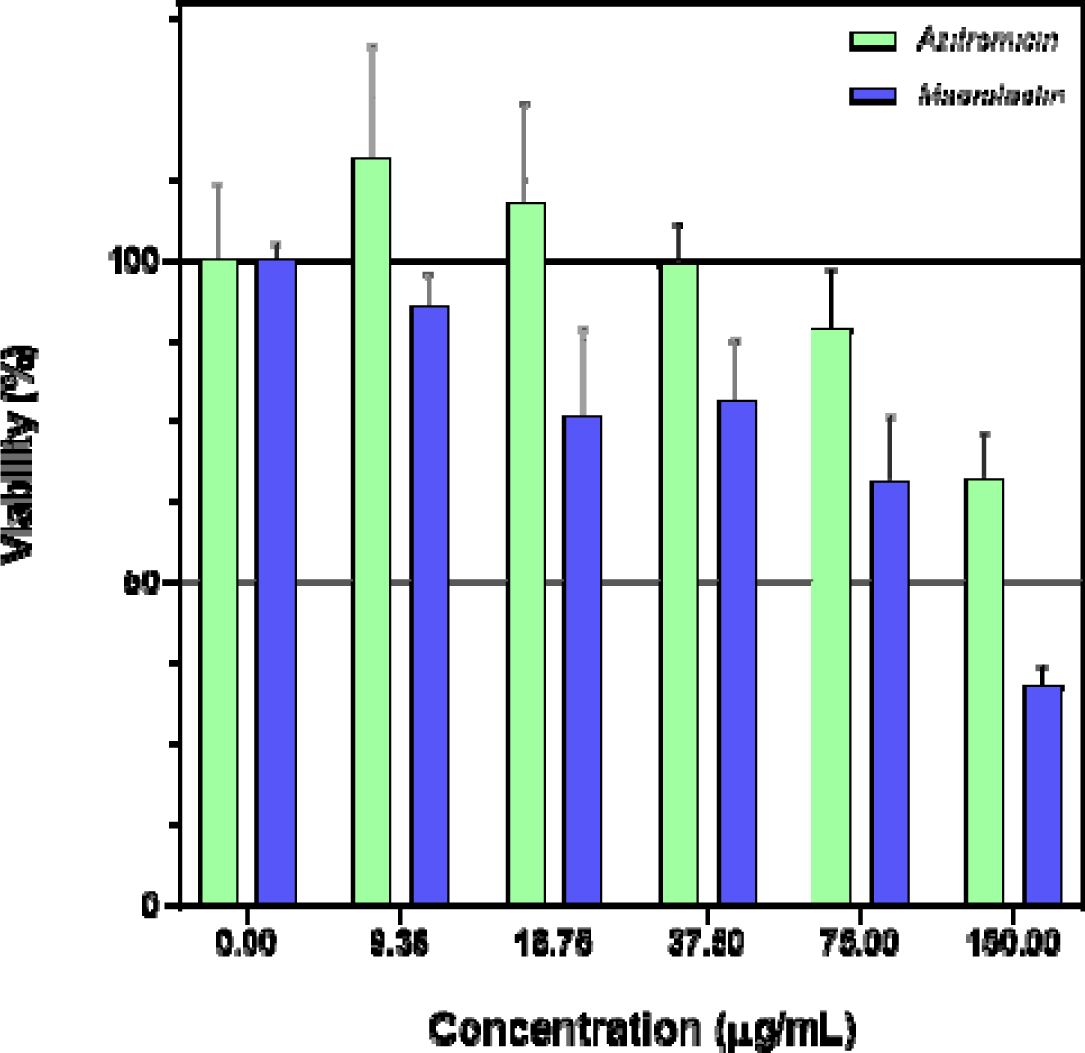
Cytotoxic property of McA in relation to keratinocyte.

Interestingly, it was not possible to dock the McA molecule with those areas where amino acid substitutions occurred. In two cases with wild and mutated proteins, docking poses were fully identical and had the same value of a docking score (-9.2 kcal/mol). From here it follows logically that the assumed mutations (notably Arg125His) create a steric hindrance the entry of McA to the identified binding site.

Further studies using molecular dynamics are needed to assess the effect of amino acid substitutions on the conformational flexibility of the EF-Tu molecule and experimental validation via structural biology methods such X-ray diffraction or cryo-EM.

### 2.6 Macrolactin A is not toxic for human keratinocytes

Discussing the potential of McA as a therapeutic agent, it is necessary to evaluate its toxicity in *in vitro* and *in vivo* models. The work of Jung et al. (2014) assessed the pharmokinetic parameters of McA after intravenous, oral, and intraperitoneal administration to mice. It was found that with intravenous administration of 50 mg/kg of McA, its concentration in the blood plasma dynamically changed from initial values of ∼ 20 μg/mL (0 min) to ∼ 0.5 μg/mL (60 min). Despite preclinical studies that indicate the low toxicity of McA, data on its cytotoxicity are sporadic. Thus, the work of Romero-Tabarez et al. (2006) revealed that McA at a concentration of approximately 30 μg/mL suppressed the proliferation of 80% of L929 mouse fibroblasts cells.

We studied cytotoxicity of McA using the MTT test according to the recommendations of ICCVAM (NIH Publication No. 07-4519, 2006). According to the results of the MTT test, the IC50 concentration for McA significantly exceeded 100 μg/mL (Figure 4), which allows us to classify this compound as low toxic (Krippendorff et al. 2007). At the same time, a study of the toxicity of McA showed a higher toxicity compared to in comparison with azithromycin, its closest analog whose toxic effect was manifested only at a concentration of 150 μg/mL.

## 3. CONCLUSION

Macrolactins are a group of natural macrolide antibiotics, comprising at least 33 structural variants (Xiao and Li, 2020), among which McA is the most studied. The biological activity of McA has been identified against both prokaryotic (antibacterial) and eukaryotic (anticancer, anti-inflammatory) organisms (Nagao et al. 2001; Yan et al. 2016; Jin et al. 2017). Studies describing the antibacterial properties of McA consist of diverse data obtained over different years using various methodologies.

There are several studies proposing hypotheses about the mechanism of the antimicrobial action of McA and its 7-O-malonyl derivative. The use of the purified McA fraction made it possible to visualize impaired cell division and separation of daughter cells, as well as incomplete formation of septa in both Gram-positive and Gram-negative cells (Chen et al. 2019). Similar observations were reported by Chen et al. (2021), where transmission electron microscopy demonstrated the inhibition of cell division in *Agrobacterium tumefaciens* C58 treated with a culture medium containing macrolactins. Apparently, macrolactins somehow impact cell division processes.

In our study, the results of inhibitory concentration assessments revealed that among both Gram-positive and Gram-negative bacteria, there are species and strains susceptible to the action of McA, as well as those that are not sensitive. Analyzing all these data, one can hypothesize that the action of McA on bacteria is independent of the type (Gram-positive or Gram-negative) of their cell wall. This fact indicates that the main target of McA in bacterial cells is unlikely biosynthesis processes of the cell wall or its integrity. In this regard, the signs of cell shape disruption observed in some studies upon exposure to McA should probably be explained by other mechanisms.

Our developments in biosensors for screening and differentiating antibiotics based on their mechanism of action, allowed us to determine that McA acts as an inhibitor of the protein biosynthesis in bacterial cells. The use of *in vitro* cell-free models demonstrated that McA indeed suppresses the protein synthesis process, with inhibitory concentrations (IC50) being lower in the case of the *E. coli* model compared to *S. aureus.* A more detailed analysis using toe-printing assay not only confirmed these results but also determined that McA induces ribosome stalling at the first translation step. To further detail the mechanism identified, involving ribosome stalling, we obtained the McA-resistant clone and analyzed its genetic characteristics that set it apart from the original sensitive strain. It turned out that *B. pumilus* McA^R^ has mutations in the gene encoding elongation factor Tu. Indirect confirmation that we are on the right track comes from the results of a proteomic analysis conducted in the study by Romero-Tabarez (2004), where EF-Tu has expressed significantly more actively under sub-inhibitory exposure to 7-O-malonyl macrolactin A.

Among antibiotics, there is a group of substances named elfamycins, consisting of different chemical entities united by the antimicrobial target – EF-Tu (Prezioso et al., 2017). EF-Tu is a member of the G protein family, which comprises GTPase enzymes binding guanosine nucleotides (GTP and GDP) and possessing the inherent capability to hydrolyze GTP to GDP. The complete structure of EF-Tu is composed of three domains, with Domain 1, also known as the G domain, playing a significant role in EF-Tu’s GTPase activity.

Kirromycin is an elfamycin, whose mechanism of activity hase been extensively studied. Structures of ribosome-bound EF-Tu with GDP and kirromycin have been obtained by x-ray crystallography and cryogenic electron microscopy, while molecular dynamics simulations showed that this antibiotic binded to the D1-D3 interface EF-Tu and thereby sterically hindered the D1 rotation (Schmeing et al., 2009; Fischer et al., 2015; Warias et al. 2019). We conducted molecular modeling of amino acid substitutions in the native structure of EF-Tu (*B. pumilus*) and identified that the substitutions Arg125His and Val197Ala are located in the kirromycin binding pocket in Domain 1. The existing list of mutations determining EF-Tu resistance to kirromycin (Abdulkarim et al. 1994) includes substitutions in amino acid positions (Gln124Arg, Tyr160Asn) close to those revealed in our study. However, the applied molecular docking approach did not confirm that MA interacts with these amino acids. We understand that the approach used is preliminary and instrumental study is needed to resolve the structure of McA-EF-Tu complex.

Analyzing existing works dedicated to elfamycins and their interaction with EF-Tu, it is possible to explain an abnormal cell shape of bacterial cells when exposed to macrolactins (Chen et al., 2019; Chen et al. 2020). In 2010, Defeu Soufo et al. discovered that MreB and EF-Tu interact within *B. subtilis* cells. EF-Tu is associated with the bacterial cytoskeleton and plays a crucial role in maintaining MreB cytoskeletal elements. They observed that a reduction in cellular EF-Tu levels affects cell shape. Furthermore, the addition of kirromycin to exponentially growing *B. subtilis* cells arrested cell growth.

The differences in IC50 for cell-free models of *E. coli* and *S. aureus* under McA exposure can also be elucidated by considering the results of elfamycin studies. Hall’s et al. (1991) study found that the GTPase activity of EF-Tu purified from *S. aureus* was 1000-fold less sensitive to kirromycin analogues compared to EF-Tu purified from *E. coli*.

In conclusion, we note that the obtained data will be useful for rethinking the existing data on the biological effects of MacA, for example, on the effect on the soil microbial community (Yuan et al., 2016), antiviral (Bharadwaj et al. 2021) or antitumor activities. As an antibiotic, McA’s activity is weak; however, as an inhibitor of protein biosynthesis, McA can alter the metaproteome of bacterial communities in their natural habitat. Undoubtedly, further detailed studies of McA will reveal the functional significance of this antibiotic in the life of microorganisms and how it can be used for the benefit of humans.

## 4. MATERIAL AND METHODS

### 4.1 Purification of macrolactin A

Previously, we described the *Bacillus velezensis* strain X-Bio-1, in whose genome clusters of biosynthetic genes controlling the biosynthesis of polyketide antibiotics, including a homologue of macrolactin, were discovered (Kravchenko et al. 2020). Cultivation of the strain was carried out in a 1.0-liter conical flask in a medium similar to LB broth (per liter of medium, gr.: tryptone - 10, yeast extract - 5, NaCl - 2) for 48 hours. Bacterial cells were separated from the culture liquid by centrifugation and filtration through 0.22 µm membranes (VWR, USA). Next, the antimicrobial component of the metabolites was extracted from the culture medium with ethyl acetate. The resulting extract was evaporated and redissolved in dimethyl sulfoxide (DMSO). Purification to a degree of > 90% was carried out using reverse-phase HPLC on a Luna C18 analytical column (4.6 × 250 mm, 130 Å, 5 µm) (Phenomenex, USA). We used buffer A (HPLC-grade water, pH 7.5) and with a linear gradient of buffer B (80% HPLC-grade acetonitrile in buffer A), detection was carried out at 220 nm.

### 4.2 Spectrometric analysis of macrolactin A

#### Liquid Chromatography-Mass Spectrometry

Liquid chromatography-mass spectrometry was performed using a UPLC/MS/MS system consisting of an Acquity UPLC chromatograph (Waters Corporation, Milford, MA, USA) and a TQD quadrupole mass-spectrometer (Waters Corporation, Milford, MA, USA) with the registration of positive ions using the ESI MS method with the Acquity BEH column C18 (1.7 microns, 50×2.1 mm, Waters Corporation, Milford, MA, USA), flow rate 0.5 mL/min, 35 °C, and elution with a gradient of 5-100% CH_3_CN in 20 mM of HCOOH for 4 min.

#### NMR analysis of macrolactin A

For NMR spectroscopy experiments the lyophilized sample were dissolved in 99.95% MeOH-d4. All 1D and 2D NMR experiments were performed with a Bruker Avance III HD 700 MHz spectrometer equipped with a QCI cryoprobe at 298 K. Assignments were made by using 1H, 13C 1D experiments and 2D 1H-1H COSY, 1H-13C HSQC and HMBC experiments. All pulse sequences were used as provided by Bruker and processing was performed using Bruker Topspin software (v. 3.6.3).

### 4.3 Determination of minimum inhibitory and bactericidal concentrations

Minimal inhibitory concentration (MIC) was determined following the Clinical and Laboratory Standards Institute (CLSI) guidelines for broth microdilution MIC assay. The bacteria (10^5^ CFU/mL) were incubated in a 96-well microtiter plate (Eppendorf, Germany) containing Mueller–Hinton II broth (MHB; Becton-Dickinson, Sparks, MD, US) with two-fold dilutions of the antibiotics. The result was evaluated after 24 h of cultivation at 37°C. The bacterial growth was assessed by scanning the absorbance at 620 nm using a spectrophotometer Multiscan GO (Thermo Scientific, Waltham, MA, US). The wells without visible bacterial growth were plated on MHB-agar and incubated. The antimicrobial activity was expressed by the MIC and the minimal bactericidal concentration (MBC), which were defined as the lowest antibiotic dose at which no visible growth in medium and growth on agar-plate were detected, respectively.

### 4.4 Time-kill assay

Time-kill study was carried out based on guideline M26-A of the CLSI [13], using non-treated polystyrene 96-well plates (Eppendorf, Hamburg, Germany). The kill kinetics of McA against bacteria were tested by incubating theme in the medium with antibiotic taken for the 1/2 MIC, 1MIC and 2MIC. Viable cell counts were determined after 0, 1, 2, 4, 6, 8, and 24 h of incubation at 37 °C by plating the serially diluted samples onto MHB-agar plates. Bactericidal activity was defined as a ≥3-log10 CFU/mL decrease, in comparison with the baseline, after 24 h of incubation.

### 4.5 Evaluation of protein biosynthesis inhibition using a reporter strain *E. coli* JW5503 Δ*tolC* pDualrep2

The “pDualrep2” system was used to evaluate the mechanism of antimicrobial action of strain extracts with antimicrobial activity. This system based on hypersensitive stain JW5503 (Δ*tolC*) (Baba et al. 2006) transformed with “pDualrep2” plasmid (Osterman et al., 2016). This system allows to sort out suppressors of protein synthesis or SOS-response induces. The biosensor has been widely used in studies investigating analogs of conventional antibiotics with improved antimicrobial activity (Tevyashov et al., 2019) or the mechanism of action of novel antibiotics (Metelev et al., 2017).

In short, 100 µL of cultural broth was added in wells in an agar plate containing the reporter strain *E. coli* JW5503 and incubated overnight at 37 °C. The agar plate was scanned using the ChemiDoc Imaging System (Bio-Rad Laboratory, US). This system consisted of two channels, “Cy3-blot” (553/574 nm, green pseudocolor) fluorescent for Turbo red fluorescent protein (TurboRFP) and “Cy5-blot” (588/633 nm, red pseudocolor) for Katushka2S fluorescence. Translation inhibitors triggered the induction of Katushka2S expression, while TurboRFP was upregulated by DNA damage-induced SOS response.

### 4.6 Inhibition assays of in cell-free translation systems

*The E.coli model.* The inhibition of firefly luciferase synthesis in cell-free translation systems by the studied compound was tested with an *E. coli* S30 Extract System for Linear Templates (Promega, USA). The reactions were carried out in 5 μL aliquots containing 2 μL of S30 Premix, 1.5 μL of S30 extract E. coli, 0.5 μL of amino acid mixture (each in 1mM), 0.5 μL of mRNA (Fluc 100 ng/UL), 0.1 μL of Ribolock (RNAze inhibitor) and 0.5 μL of tested compound. The reaction mixture (except for mRNA) was pre-mixed on ice and then incubated at room temperature for 5 min to give the antibiotic ample time to bind to the ribosome be-fore the initiator complex assembly; the mixture was then returned on ice, and the template was added. Translation was carried out for 20 min at 37°C. The activity of in vitro synthesized luciferase was assessed using 5 μL of the substrate from the Steady-Glo Luciferase Assay System (Promega, US). The signal was detected on plate reader Victor X5 2030 (Perkin Elmer, US).

*The S.aureus model.* To analyze the influence of McA on protein synthesis in Gram-positive bacteria, we used the coupled transcription-translation cell-free system, based on *S. aureus* (strain 4220) cell extract (S30) and sfGFP as a fluorescent reporter (Murray et al. 2001). The protocols of extract and reaction mix preparations, the conditions of kinetic experiments were taken from the study described before (Fatkhullin et al., 2022). The concentrations of 14 mM for Mg^2+^, 370 mM for K^+^, and of 35% for extract were used as optimal. Kinetic measurements were performed in presence of macrolactin A (dissolved in DMSO) of different concentrations in reaction mix (0.01, 0.05, 0.1, 1, 10, 50, 100, 200 μg/ml). The variants with the addition of a DMSO and without the addition of a reporter plasmid were used as positive and negative controls, respectively. The sfGFP fluorescence had been measured every 5 minutes during 2 hours (485 nm (excitation) and 510 nm (emission). Erythromycin taken for 25 μg/mL was used as a positive control. The kinetic measurements of sfGFP fluorescence in a cell-free reaction mix were performed in 384-well Greiner black plates by a Varioskan LUX Multimode Microplate Reader (Thermo Fisher Scientific, US).

### 4.7 Toe-printing analysis

The toe-printing assay was conducted according to the protocol described by Orelle et al. (2013). At the first stage, the primers were labeled with [γ-32P] ATP polynucleotide kinase (ThermoFisher, US) according to the manufacturer’s protocol. Next, in vitro translation of the short-model mRNA was performed using a PURExpress® In VitroProtein Synthesis Kit (New England Biolabs, US). The reaction mixture (volume, 5 μL) contained 2 μL of solution A, 1 μL of solution B, 0.2 μL of RiboLock (ThermoFisher), 0.5 μL of the test compound, 0.5 μL of DNA template (0.2 mmol/μL), and 0.5 μL of the radiolabeled primer.

The mixture was incubated at 37°C for 20 min, and 1 μL of the reverse transcrip-tion mix from the Titan One Tube RT-PCR System kit (Roche, Switzerland) was added. Reverse tran-scription was conducted for 15 min at 37°C. The re-action was stopped by adding 1 μL of 10 M NaOH, followed by incubation at 37°C for 15 min. The neu-tralization was performed by adding 1 μL of 10 N HCl. Next, 200 μL of the resuspension buffer as added.The resulting samples were purified using a QIAquick PCR purification kit (Qiagen, Germany).The sequence mixtures were prepared using a USB® Thermo Sequenase Cycle Sequencing Kit (Affymetrix, USA) according to the manufacturer’s protocol. Electrophoresis was carried out in 6% polyacryla-mide gel (60 × 40 × 0.03 cm) containing 19% acryla-mide, 1% N,N’- methylenebisacrylamide, and 7 M urea in TBE buffer for 2–3 h.

The specimens and products of the sequencing reactions (2 and 1.5 μL, respectively) were applied onto the gel. The gel was transferred onto 3-mm paper, dried, and exposed to a sensory screen for 18 h. The screen was scanned using a Typhoon FLA 9500 Biomolecular Imager (GE Healthcare, US). The ermCL template for this experiment was obtained by PCR amplification using a Taq-DNA-polymerase kit (ThermoFisher, US), according to the standard protocol. The template sequence described in Table S2.

### 4.8 Serial passage experiment

Overnight culture of *Bacillus pumilus* UTMN was inoculated into LB broth containing McA in twofold serial dilutions as described (Poshvina et al. 2022). The cells were incubated at 37°C for 24 h. After incubation, bacterial cells growing at subinhibitory concentrations of McA were used for subsequent passaging. The procedure was repeated for 20 times. The MIC increased from 4 to 38 μg/mL after 20 passages (Figure S2).

### 4.9 DNA extraction, genome sequencing and annotation

Genomic DNA was isolated from overnight broth culture using the Quick-DNA Fungal/Bacterial Miniprep kit (ZymoResearch, USA) according to the manufacturer’s protocol. The concentration of the DNA was estimated by utilizing the Qubit 4.0 fluorimeter manufactured (Thermo Fisher Scientific, Germany). The quality of the DNA was assessed using the NanoPhotometer N120 (Implen, USA) spectrophotometer. Genomic libraries were prepared using the NEBNext Ultra II FS library preparation kit (New England Biolabs, USA) according to the manufacturer’s instructions for MiSeq (Illumina, US) sequencing.

The whole genome sequencing was carried out on the MiSeq instrument (Illumina, USA) using a MiSeq reagent kit V3 with 2×250-bp reads. Quality of the reads were assessed using the FastQC program v0.11.5. The obtained reads did not contain adapter sequences. The reads were filtered based on quality (Q=20) and length (removing the first and last 10 nucleotides, as well as all discarding reads shorter than 50 bp) using the Trimmomatic v0.39 (Bolger et al., 2014). Genome assembly was performed using the Unicycler program v0.4.9b (Wick et al., 2017), including assembly using the SPAdes genome assembler v3.15.3. The resulting contigs were analyzed using the Quast program (Gurevich et al., 2013) to assess the assembly quality. Contigs shorter than 200 base pairs were excluded from further analysis. Identification of SNP was performed using the Snippy software (https://github.com/tseemann/snippy). The fastq clean reads of resistant strain *B. pumilus* McA^R^ were aligned against complete genome of wild type strain *B. pumilus*.

The assembled genome sequence of *B. pumilus* McA^S^ and *B. pumilus* McA^R^ were deposited in GenBank under accession no. JAWZXI000000000 and JAWZXH000000000, respectively. The Illumina MiSeq raw reads were deposited in the NCBI Sequence Read Archive (SRA) under accession no. SRR26911262 and SRR26911261 (BioProject no. PRJNA1041609; BioSample no. SAMN38289444 and SAMN38289364).

### 4.10 Homology modeling and molecular docking

Because experimental structural data currently is not available, we performed a homology modeling of *Bacillus pumilus* EF-Tu. High-quality X-ray structure of *Thermus thermophilus* strain HB8 EF-Tu in an active (GTP-binding) conformation was used for template (PDB ID: 2C78) (Parmeggiani et al. 2006). Reference sequence of studied protein obtained from the UniProt database (entry: P0A6N3). Wild and McA resistant EF-Tu proteins were modeled in the SWISS-MODEL (Waterhouse et al 2018). We used Smina software (version Nov 9 2017) for molecular docking (Koes et al. 2013). McA structure was obtained from the PubChem database (PubChem CID: 6451096). Protein structures were prepared in the OpenBabel (version 3.1.1): protonation and adding charges (O’Boyle et al. 2011). GRID-box size was 40×40×40 Å with a center in the co-crystallized ligand PUL (pulvomycin). Visualization and molecular structures manipulations were conducted in the PyMol (Schrödinger, Inc; version 2.5.0; Open-Source Build), 2D interaction map was prepared Maestro (Schrödinger, Inc; version 13.0.137; Academic license).

### 4.11 Cytotoxic activity of macrolactin A against human cells

The normal immortalized keratinocyte line HaCaT from the DKFZ collection (CLS Cell Lines Service, 300493) was used as a model object to determine cytotoxicity. After thawing, the cells were passaged twice in DMEM/F12 culture medium (PanEco, Russia) containing 10% FBS (Sigma, USA) with the addition of 1% GlutaMAX (Gibco, USA) and penicillin/streptomycin solution (100 units/mL and 100 μg/mL respectively, PanEco, Russia). Cells were grown in culture flasks with an area of 75 cm2. The medium was replaced with fresh one every second day of cultivation. Upon reaching the exponential phase of division, the cells were collected by trypsinization and then seeded into the wells of a 96-well plate at a rate of 2.0 × 103 cells per well 24 hours before adding the test substances.

The cytotoxicity of the studied substances was assessed using the MTT test. The test substances were dissolved in DMSO, after which serial dilutions of the resulting solutions were prepared in the culture medium with a dilution factor of 2 so that the final DMSO concentration did not exceed 0.5%. The toxicity of the analyzed substances was assessed in the concentration range of 9.375-150 μg/mL. 24 hours after cell seeding, the culture medium was replaced with one containing the test substances. After the exposure time with the test substances had expired, the culture medium was replaced with one containing MTT (1 mg/mL) and incubated under standard conditions for three hours, after which the cells were washed with DPBS solution and lysed in DMSO. Optical density was measured using a Bio-Rad iMark tablet spectrophotometer at 595 nm). In control groups, the culture medium was replaced with one containing 0.5% DMSO.

## Author Contributions

Conceptualization ASV; investigation ASV, DAL, DSD, KSU, DVP, AAN, ANI, NSG, AT, AGB, EAK, DDR, NGL, AVT; funding acquisition ASV, DVP, DAL, AT, KSU; supervision ASV, IAO, PVS, KSU, ALR; writing original draft ASV, DAL, KSU, AVT; writing review and editing ASV, DAL, KSU. All authors have read and agreed to the published version of the manuscript.

## Data Availability Statement

The Illumina MiSeq raw reads were deposited in the NCBI Sequence Read Archive (SRA) under accession no. SRR26911262 and SRR26911261 (BioProject no. PRJNA1041609; BioSample no. SAMN38289444 and SAMN38289364).

## Acknowledgments

We are grateful to V.N. Tashlitsky (Lomonosov Moscow State University) and Irina V. Palamarchuk (University of Tyumen) for the LC-MS analysis.

## Supplementary files

**Figure S1.**
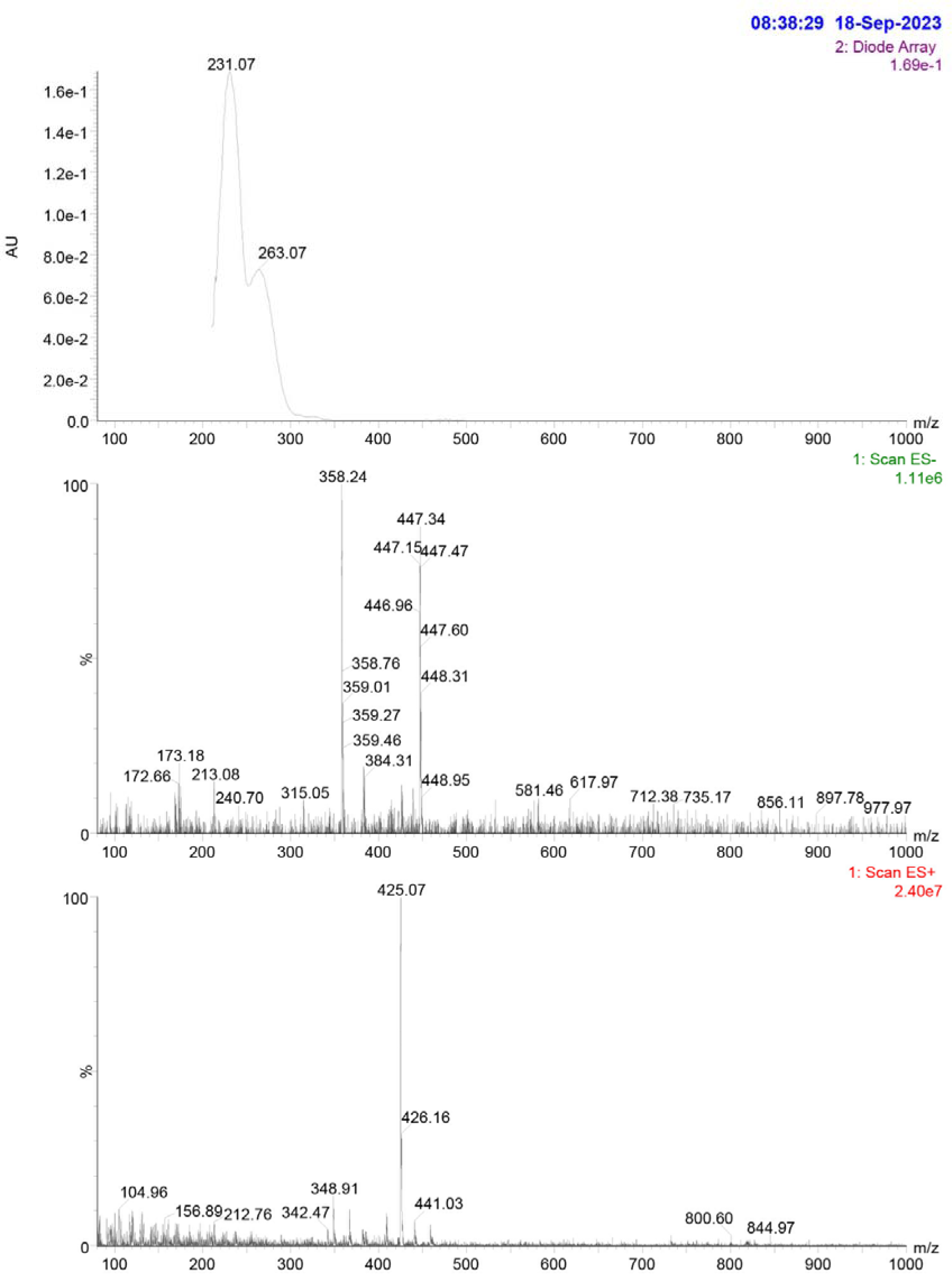
UV-spectrum of antimicrobial metabolite of *B. velezensis* X-bio-1 (a); and mass-spectraа obtained in negative (b) and positive ESI regimes (c).

**Figure S1.**
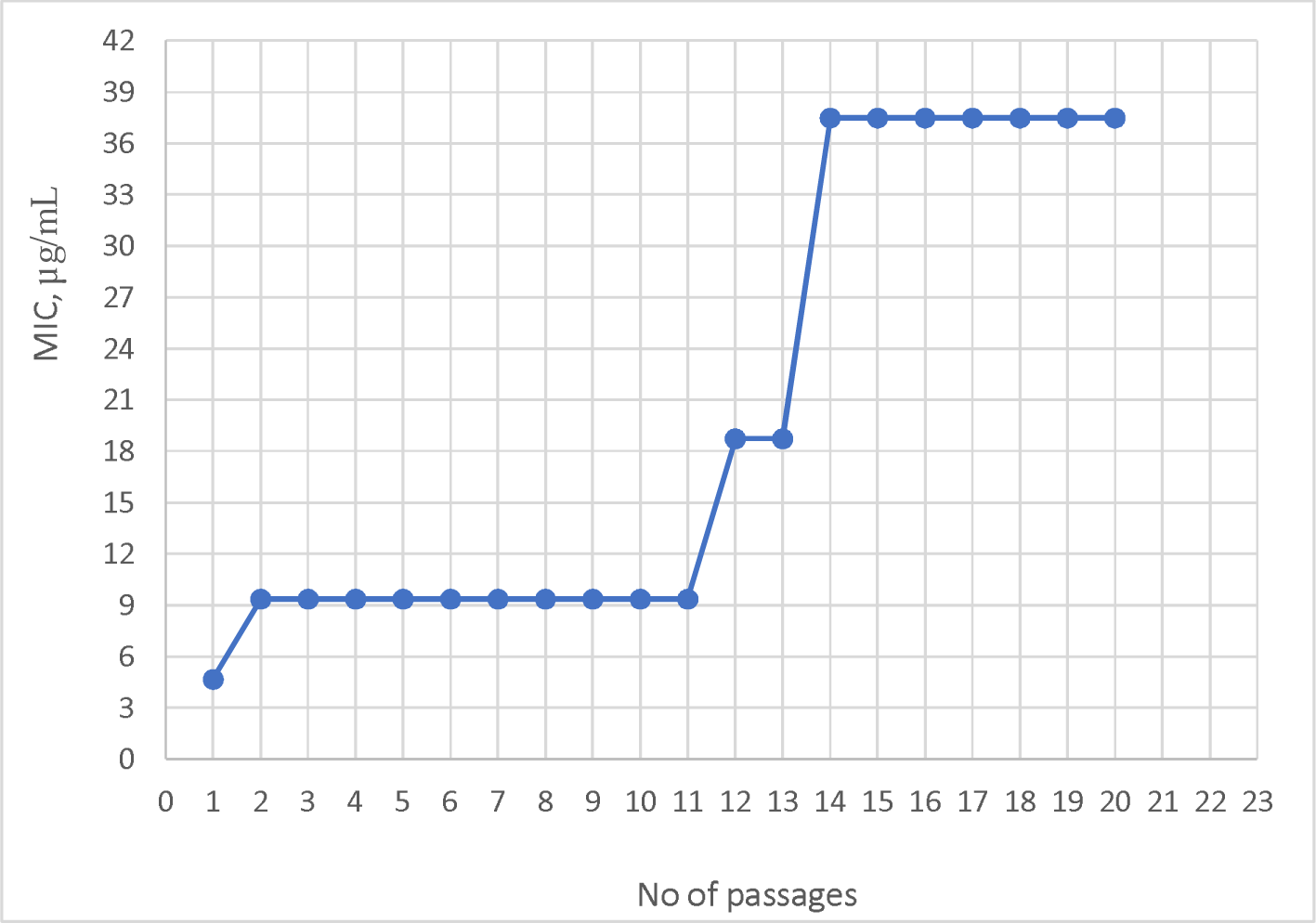
Dynamics of development of *Bacillus pumilus* UTMN resistance to macrolactin A

## References

1. Abdulkarim F, Liljas L, Hughes D. Mutations to kirromycin resistance occur in the interface of domains I and III of EF-Tu·GTP. FEBS Lett 1994;352:118–22. 10.1016/0014-5793(94)00937-6.

2. Baba T, Ara T, Hasegawa M, Takai Y, Okumura Y, Baba M, et al. Construction of Escherichia coli K-12 in-frame, single-gene knockout mutants: The Keio collection. Mol Syst Biol 2006;2. 10.1038/msb4100050.

3. Barry AL, A. W, Nadler MDH, Reller PDLB, M.D. Christine C. Sanders, Ph.D. Jana M. Swenson MMS. M26-A Methods for Determining Bactericidal Activity of Antimicrobial Agents; Approved Guideline This document provides procedures for determining the lethal activity of antimicrobial agents. Clin Lab Stand Inst 1999;19:1–14.

4. Belanger CR, Lee AHY, Pletzer D, Dhillon BK, Falsafi R, Hancock REW. Identification of novel targets of azithromycin activity against Pseudomonas aeruginosa grown in physiologically relevant media. Proc Natl Acad Sci U S A 2020;117:33519–29. 10.1073/pnas.2007626117.

5. Bolger AM, Lohse M, Usadel B. Trimmomatic: A flexible trimmer for Illumina sequence data. Bioinformatics 2014;30:2114–20. 10.1093/bioinformatics/btu170.

6. Bharadwaj KK, Sarkar T, Ghosh A, Baishya D, Rabha B, Panda MK, Nelson BR, John AB, Sheikh HI, Dash BP, Edinur HA, Pati S. Macrolactin A as a Novel Inhibitory Agent for SARS-CoV-2 M^pro^: Bioinformatics Approach. Appl Biochem Biotechnol. 2021 Oct;193(10):3371–3394. doi: 10.1007/s12010-021-03608-7.

7. Chen J, Liu T, Wei M, Zhu Z, Liu W, Zhang Z. Macrolactin a is the key antibacterial substance of Bacillus amyloliquefaciens D2WM against the pathogen Dickeya chrysanthemi. Eur J Plant Pathol 2019;155:393–404. 10.1007/s10658-019-01774-3.

8. Chen L, Wang X, Liu Y. Contribution of macrolactin in Bacillus velezensis CLA178 to the antagonistic activities against Agrobacterium tumefaciens C58. Arch Microbiol 2021;203:1743–52. 10.1007/s00203-020-02141-1.

9. Elkashif A, Seleem MN. Investigation of auranofin and gold-containing analogues antibacterial activity against multidrug-resistant Neisseria gonorrhoeae. Sci Rep 2020;10:1–9. 10.1038/s41598-020-62696-3.

10. Fatkhullin B, Golubev A, Garaeva N, Validov S, Gabdulkhakov A, Yusupov M. Y98 Mutation Leads to the Loss of RsfS Anti-Association Activity in Staphylococcus aureus. Int J Mol Sci 2022;23. 10.3390/ijms231810931.

11. Fischer N, Neumann P, Konevega AL, Bock L V., Ficner R, Rodnina M V., et al. Structure of the E. coli ribosome-EF-Tu complex at <3 Å resolution by Cs-corrected cryo-EM. Nature 2015;520:567–70. 10.1038/nature14275.

12. Gustafson K, Roman M, Fenical W. The macrolactins, a novel class of antiviral and cytotoxic macrolides from a deep-sea marine bacterium. Journal of the American Chemical Society 1989;111:7519–7524.

13. Gurevich A, Saveliev V, Vyahhi N, Tesler G. QUAST: Quality assessment tool for genome assemblies. Bioinformatics 2013;29:1072–5. 10.1093/bioinformatics/btt086.

14. Hall CC, Watkins JD, Georgopapadakou NH. Comparison of the Tu elongation factors from Staphylococcus aureus and Escherichia coli: Possible basis for elfamycin insensitivity. Antimicrob Agents Chemother 1991;35:2366–70. 10.1128/AAC.35.11.2366.

15. Htwe Maung CE, Choub V, Cho JY, Kim KY. Control of the bacterial soft rot pathogen, Pectobacterium carotovorum by Bacillus velezensis CE 100 in cucumber. Microb Pathog 2022;173. 10.1016/j.micpath.2022.105807.

16. Han JS, Cheng JH, Yoon TM, Song J, Rajkarnikar A, Kim WG, et al. Biological control agent of common scab disease by antagonistic strain Bacillus sp. sunhua. J Appl Microbiol 2005;99:213–21. 10.1111/j.1365-2672.2005.02614.x.

17. Jin J, Choi SH, Lee JE, Joo JD, Han JH, Park SY, et al. Antitumor activity of 7-O-succinyl macrolactin A tromethamine salt in the mouse glioma model. Oncol Lett 2017;13:3767–73. 10.3892/ol.2017.5918.

18. Jung JW, Kim JM, Kwon MH, Kim DH, Kang HE. Pharmacokinetics of macrolactin A and 7-O-succinyl macrolactin A in mice. Xenobiotica 2014;44:547–54. 10.3109/00498254.2013.861542.

19. Koes DR, Baumgartner MP, Camacho CJ. Lessons learned in empirical scoring with smina from the CSAR 2011 benchmarking exercise. J Chem Inf Model. 2013 Aug 26;53(8):1893–904. doi: 10.1021/ci300604z.

20. Krippendorff BF, Lienau P, Reichel A, Huisinga W. Optimizing classification of drug-drug interaction potential for CYP450 isoenzyme inhibition assays in early drug discovery. J Biomol Screen 2007;12:92–9. 10.1177/1087057106295897.

21. Kravchenko S V., Poshvina D V., Vasilchenko A V., Vasilchenko AS. Draft Genome Sequence of the Multiple Antibiotic Producer Bacillus velezensis X-BIO-1. Microbiol Resour Announc 2020;9:19–20. 10.1128/mra.01319-20.

22. Kim DH, Kim HK, Kim KM, Kim CK, Jeong MH, Ko CY, et al. Antibacterial activities of macrolactin a and 7-O-succinyl macrolactin a from Bacillus polyfermenticus KJS-2 against vancomycin-resistant enterococci and methicillin-resistant Staphylococcus aureus. Arch Pharm Res 2011;34:147–52. 10.1007/s12272-011-0117-0.

23. Li W, Tang XX, Yan X, Wu Z, Yi ZW, Fang MJ, et al. A new macrolactin antibiotic from deep sea-derived bacteria Bacillus subtilis B5. Nat Prod Res 2016;30:2777–82. 10.1080/14786419.2016.1155576.

24. Lewis K. The Science of Antibiotic Discovery. Cell 2020;181:29–45. 10.1016/j.cell.2020.02.056.

25. Lemaire S, Van Bambeke F, Tulkens PM. Cellular accumulation and pharmacodynamic evaluation of the intracellular activity of CEM-101, a novel fluoroketolide, against Staphylococcus aureus, Listeria monocytogenes, and Legionella pneumophila in human THP-1 macrophages. Antimicrob Agents Chemother 2009;53:3734–43. 10.1128/AAC.00203-09.

26. Myers AG, Clark RB. Discovery of Macrolide Antibiotics Effective against Multi-Drug Resistant Gram-Negative Pathogens. Acc Chem Res 2021;54:1635–45. 10.1021/acs.accounts.1c00020.

27. Murray RW, Melchior EP, Hagadorn JC, Marotti KR. Staphylococcus aureus cell extract transcription-translation assay: Firefly luciferase reporter system for evaluating protein translation inhibitors. Antimicrob Agents Chemother 2001;45:1900–4. 10.1128/AAC.45.6.1900-1904.2001.

28. Metelev M, Osterman IA, Ghilarov D, Khabibullina NF, Yakimov A, Shabalin K, et al. Klebsazolicin inhibits 70S ribosome by obstructing the peptide exit tunnel. Nat Chem Biol 2017;13:1129–36. 10.1038/nchembio.2462.

29. Martin Schmeing T, Voorhees RM, Kelley AC, Gao YG, Murphy IV F V., Weir JR, et al. The crystal structure of the ribosome bound to EF-Tu and aminoacyl-tRNA. Science (80-) 2009;326:688-94. 10.1126/science.1179700.

30. Nagao T, Adachi K, Sakai M, Nishijima M, Sano H. Novel macrolactins as antibiotic lactones from a marine bacterium. J Antibiot (Tokyo) 2001;54:333–9. 10.7164/antibiotics.54.333.

31. O’Boyle NM, Banck M, James CA, Morley C, Vandermeersch T, Hutchison GR. Open Babel: An open chemical toolbox. J Cheminform. 2011 Oct 7;3:33. doi: 10.1186/1758-2946-3-33.

32. Osterman IA, Komarova ES, Shiryaev DI, Korniltsev IA, Khven IM, Lukyanov DA, et al. Sorting out antibiotics’ mechanisms of action: A double fluorescent protein reporter for high-throughput screening of ribosome and DNA biosynthesis inhibitors. Antimicrob Agents Chemother 2016;60:7481–9. 10.1128/AAC.02117-16.

33. Orelle C, Carlson S, Kaushal B, Almutairi MM, Liu H, Ochabowicz A, et al. Tools for characterizing bacterial protein synthesis inhibitors. Antimicrob Agents Chemother 2013;57:5994–6004. 10.1128/AAC.01673-13.

34. Prezioso SM, Brown NE, Goldberg JB. Elfamycins: inhibitors of elongation factor-Tu. Molecular microbiology 2017;106:22–34. 10.1111/mmi.13750

35. Poshvina D V., Dilbaryan DS, Kasyanov SP, Sadykova VS, Lapchinskaya OA, Rogozhin EA, et al. Staphylococcus aureus is able to generate resistance to novel lipoglycopeptide antibiotic gausemycin A. Front Microbiol 2022;13. 10.3389/fmicb.2022.963979.

36. Parmeggiani A, Krab IM, Okamura S, Nielsen RC, Nyborg J, Nissen P. Structural basis of the action of pulvomycin and GE2270 A on elongation factor Tu. Biochemistry. 2006 Jun 6;45(22):6846–57.

37. Romero-Tabarez M, Jansen R, Sylla M, Lünsdorf H, Häußler S, Santosa DA, et al. 7-O-malonyl macrolactin A, a new macrolactin antibiotic from Bacillus subtilis active against methicillin-resistant Staphylococcus aureus, Vancomycin-resistant enterococci, and a small-colony variant of Burkholderia cepacia. Antimicrob Agents Chemother 2006;50:1701–9. 10.1128/AAC.50.5.1701-1709.2006.

38. Romero-Tabarez M. Discovery of the new antimicrobial compound 7-O-malonyl macrolactin A, Doctoral Thesis. 2004, [accessed 29 December 2023]. 10.24355/dbbs.084-200511080100-376.

39. Sohn MJ, Zheng CJ, Kim WG. Macrolactin S, a new antibacterial agent with FabG-inhibitory activity from Bacillus sp. AT28. J Antibiot (Tokyo) 2008;61:687-91. 10.1038/ja.2008.98.

40. Soufo HJD, Reimold C, Linne U, Knust T, Gescher J, Graumann PL. Bacterial translation elongation factor EF-Tu interacts and colocalizes with actin-like MreB protein. Proc Natl Acad Sci U S A 2010;107:3163–8. 10.1073/pnas.0911979107.

41. Singkham-In U, Muhummudaree N, Chatsuwan T. In vitro synergism of azithromycin combination with antibiotics against oxa-48-producing klebsiella pneumoniae clinical isolates. Antibiotics 2021;10. 10.3390/antibiotics10121551.

42. Tacconelli E, Carrara E, Savoldi A, Harbarth S, Mendelson M, Monnet DL, et al. Discovery, research, and development of new antibiotics: the WHO priority list of antibiotic-resistant bacteria and tuberculosis. Lancet Infect Dis 2018;18:318–27. 10.1016/S1473-3099(17)30753-3.i

43. Tevyashova AN, Bychkova EN, Korolev AM, Isakova EB, Mirchink EP, Osterman IA, et al. Synthesis and evaluation of biological activity for dual-acting antibiotics on the basis of azithromycin and glycopeptides. Bioorganic Med Chem Lett 2019;29:276–80. 10.1016/j.bmcl.2018.11.038.

44. Waterhouse A., Bertoni M., Bienert S., Studer G.,Tauriello G., Gumienny R., Heer F.T., de Beer T.A.P.,Rempfer C., Bordoli L., Lepore R., Schwede T., SWISS-MODEL: homology modelling of protein structures and complexes, *Nucleic Acids Research*, Volume 46, Issue W1, 2 July 2018, Pages W296–W303, 10.1093/nar/gky427

45. Warias M, Grubmüller H, Bock L V. tRNA Dissociation from EF-Tu after GTP Hydrolysis: Primary Steps and Antibiotic Inhibition. Biophys J 2020;118:151–61. 10.1016/j.bpj.2019.10.028.

46. Wick RR, Judd LM, Gorrie CL, Holt KE. Unicycler: Resolving bacterial genome assemblies from short and long sequencing reads. PLoS Comput Biol 2017;13:1–22. 10.1371/journal.pcbi.1005595.

47. Wu T, Xiao F, Li W. Macrolactins: biological activity and biosynthesis. Mar Life Sci Technol 2021;3:62-8. 10.1007/s42995-020-00068-6.

48. Yuan J, Zhao M, Li R, Huang Q, Rensing C, Raza W, et al. Antibacterial compounds- macrolactin alters the soil bacterial community and abundance of the gene encoding PKS. Front Microbiol 2016;7:1–10. 10.3389/fmicb.2016.01904.

49. Yoo JS, Zheng CJ, Lee S, Kwak JH, Kim WG. Macrolactin N, a new peptide deformylase inhibitor produced by Bacillus subtilis. Bioorganic Med Chem Lett 2006;16:4889–92. 10.1016/j.bmcl.2006.06.058.

50. Yan X, Zhou YX, Tang XX, Liu XX, Yi ZW, Fang MJ, et al. Macrolactins from marine-derived bacillus subtilis b5 bacteria as inhibitors of inducible nitric oxide and cytokines expression. Mar Drugs 2016;14:1–8. 10.3390/md14110195.

51. Zotchev SB, Stepanchikova A V., Sergeyko AP, Sobolev BN, Filimonov DA, Poroikov V V. Rational design of macrolides by virtual screening of combinatorial libraries generated through in silico manipulation of polyketide synthases. J Med Chem 2006;49:2077–87. 10.1021/jm051035i.

52. Zhang, L., Jin, M., Shi, X. et al. Macrolactin Metabolite Production by *Bacillus* sp. ZJ318 Isolated from Marine Sediment. Appl Biochem Biotechnol 194, 2581–2593 (2022). 10.1007/s12010-022-03841-8

